# Differential activation of rhodopsin triggers distinct endocytic trafficking and recycling in vivo via differential phosphorylation

**DOI:** 10.1101/2024.01.03.574057

**Authors:** Darwin Ferng, Wesley Sun, Bih-Hwa Shieh

## Abstract

Activated GPCRs are phosphorylated and internalized mostly via clathrin-mediated endocytosis (CME), which are then sorted for recycling or degradation. We investigated how differential activation of the same GPCR affects its endocytic trafficking *in vivo* using rhodopsin as a model in flies expressing mCherry-tagged rhodopsin 1 (Rh1-mC) or GFP-tagged arrestin 1 (Arr1-GFP). Upon blue light stimulation, activated Rh1 recruited Arr1-GFP to the rhabdomere, which became co-internalized and accumulated in cytoplasmic vesicles of photoreceptors. This internalization was eliminated in *shi^ts1^*mutants affecting dynamin. Moreover, it was blocked by either *rdgA* or *rdgB* mutations affecting the PIP_2_ biosynthesis. Together, the blue light-initiated internalization of Rh1 and Arr1 belongs to CME. Green light stimulation also triggered the internalization and accumulation of activated Rh1-mC in the cytoplasm but with faster kinetics. Importantly, Arr1-GFP was also recruited to the rhabdomere but not co-internalized with Rh1-mC. This endocytosis was not affected in *shi^ts1^* nor *rdgA* mutants, indicating it is not CME. We explored the fate of internalized Rh1-mC following CME and observed it remained in cytoplasmic vesicles following 30 min of dark adaptation. In contrast, in the non-CME Rh1-mC appeared readily recycled back to the rhabdomere within five min of dark treatment. This faster recycling may be regulated by rhodopsin phosphatase, RdgC. Together, we demonstrate two distinct endocytic and recycling mechanisms of Rh1 via two light stimulations. It appears that each stimulation triggers a distinct conformation leading to different phosphorylation patterns of Rh1 capable of recruiting Arr1 to rhabdomeres. However, a more stable interaction leads to the co-internalization of Arr1 that orchestrates CME. A stronger Arr1 association appears to impede the recycling of the phosphorylated Rh1 by preventing the recruitment of RdgC. We conclude that conformations of activated rhodopsin determine the downstream outputs upon phosphorylation that confers differential protein-protein interactions.

## INTRODUCTION

GPCRs are critically involved in human physiology and pathophysiology, and many are targeted for pharmacological intervention. Each GPCR is tightly regulated to ensure both the temporal and the spatial resolution of the signaling response. Prolonged activation of GPCRs results in desensitization and down regulation, which is often accompanied by internalization of activated receptors. Specifically, activated GPCRs can be phosphorylated at the C-terminus and/or the third cytoplasmic loop by distinct protein kinases including G-protein-coupled receptor kinases (GRKs) (1, 2). Phosphorylation of the receptor promotes the association with arrestin that tethers activated GPCRs for internalization and alternate signaling (3).

The molecular mechanisms of GPCR endocytosis appear highly conserved. Endocytic trafficking of activated GPCRs is regulated by members of the arrestin family (4). In vertebrates, it has been shown that both arrestin 2 (β-arrestin 1) and arrestin 3 (β-arrestin 2) orchestrate endocytosis by tethering activated GPCRs to clathrin and adaptor protein-2 (3, 4). The interaction between arrestin and the receptor may be stable which leads to the co-internalization of arrestin with the receptor. Subsequently, arrestin may regulate ubiquitination of the internalized receptor affecting recycling or lysosomal degradation of the receptor (5–8). Moreover, arrestin may tether the internalized GPCRs to alternate signaling events such as the activation of ERK1/2 (9, 10).

Most studies of GPCR endocytosis were performed in cell cultures in which receptors were transiently or stably expressed. It was reported that the internalization of GPCRs is regulated in a receptor-specific manner as observed in the two related purinergic P2Y1 and P2Y12 receptors, and the dopamine D1 and D2 receptors. Here, each receptor was phosphorylated by distinct protein kinases and sorted via clathrin-mediated endocytosis (CME) or non-CME. Specifically, both P2Y receptors were internalized via CME following phosphorylation by GRK. However, endocytosis of P2Y12 is arrestin-dependent while endocytosis of P2Y1 is not (11). In contrast, the dopamine D1 receptor is internalized via CME following phosphorylation by GRK while the D2 receptor is internalized via the dynamin-independent mechanism after phosphorylation by PKC (12). It is of great interest to investigate how GPCRs can be regulated in their native cellular environment as the diversity of GRK and arrestin isoforms are likely to impact the endocytic mechanisms. Moreover, it remains to be explored how different agonists affect the trafficking of the same receptor, and how different endocytic mechanisms modulate the fate of internalized receptors.

Here we employed *Drosophila* rhodopsin as a model GPCR for insights into endocytic trafficking in its native cellular environment. Rhodopsin is critical for visual signaling and its activity is fine-tuned by several mechanisms. In the compound eye, the major rhodopsin Rh1 expressed in R1-6 photoreceptors is regulated by two distinct arrestins, arrestin 1 (Arr1) and arrestin 2 (Arr2) (13, 14). Both serve to terminate the signaling of activated Rh1 in adults (15) while Arr1 is also involved in the endocytosis of Rh1 during the development of pupal photoreceptors (16). Rh1 can be activated by a range of light with the absorption maximum centered on the blue light (480 nm) (17, 18). Thus, blue light (460-500 nm) maximally activates Rh1 similar to the effect of a full agonist on its cognate GPCR. Rh1 also can be activated by the green light (530-560 nm), possibly acting like a partial agonist, to achieve an active conformation. We explored how different conformations of activated Rh1 elicited by different wavelengths of light regulate its endocytic trafficking.

To visualize the real-time endocytosis in live photoreceptors, we employed fluorophore-tagged Rh1 (Rh1-mC) and/or Arr1 (Arr1-GFP). We previously reported that blue light stimulation triggered the internalization of Arr1-GFP. We further explored the mechanism and demonstrated the critical role of dynamin (19) and the PIP_2_ content (20, 21) in promoting internalization, supporting the notion that the blue light-induced endocytosis of Arr1-GFP belongs to CME (22). We also show that stimulation by the green light leads to internalization of Rh1-mC in pupal photoreceptors. Significantly, this endocytosis is independent of CME. Here Arr1 is recruited to the rhabdomere but is not co-internalized. Our findings also suggest that a transient interaction with Arr1 promotes faster recycling of the internalized Rh1 by facilitating dephosphorylation by rhodopsin phosphatase.

## MATERIALS AND METHODS

### Drosophila Stocks

The following fly mutants including *arr1^1^, arr2^3^, ninaC^3^, norpA^P24^, rdgA^3^*, *rdgB^KS222^,* and *shi^ts1^* were obtained from Bloomington Drosophila Stock Center (BDSC). Standard crosses were made to introduce chromosomes containing Rh1-mC and/or Arr1-GFP transgenes into various genetic backgrounds. Mutants were verified by Western blotting or behavior phenotypes.

### Fly Handling for Microscopy

Adult flies or pupae were sorted and manipulated under a dissecting microscope. This was done under ambient room light (300 lux) for less than three minutes to avoid pre-emptive rhodopsin endocytosis. Adult flies were anesthetized with CO_2_, sorted, and immobilized with clay that was placed on a Petri dish. Both males and females were used, as they served as mutants and controls respectively in F1 offspring of the X-linked mutations (*norpA*, *rdgA*, *rdgB*, and *shi*). For examining the light-dependent endocytosis in pupal photoreceptors, late-stage pupae (>p13 or >80% pupal development) (23) in the white-eyed background were used. Pupae were removed from vials and placed on double-sided tape in a glass microscope slide. The puparium covering the eye was carefully removed and the pupae were then immobilized in clay and placed on a Petri dish. The sex of the pupae was determined by the presence of the sex comb in males. Light intensity was measured using a handheld light meter.

### Manipulation of *shi^1ts^* mutants

The *shi^ts1^ (shibire)* mutants were maintained at 22°C and verified by their paralytic phenotype in adult flies when shifting to 30°C, the restrictive temperature in which Shibire/dynamin becomes inactivated. To analyze the phenotype of the mutants, the temperature of the microscope room was set to 30°C.

### Fluorescence Microscopy

Adult and pupal eyes were examined using an upright Olympus AX70 microscope equipped with a 10X lens for detecting dpp (deep pseudopupil) or a 40X water immersion lens (LUMPLFL 40X) for examining individual ommatidia. Image acquisition was performed at 100X magnification for dpp and 400X or 1000X for rhabdomeres/ommatidia using IPLab image acquisition software (BioVision Technologies, Exton, PA, USA) and the Retiga camera from QImaging (Surrey, BC, Canada). Exposure time was made constant throughout the experiment based on the brightest signal in the control group. Multiple flies (n ≥ 5) were analyzed.

### Preparation of dissociated photoreceptors

Pupal retinas were isolated and placed in a glass slide with a drop of 1X PBS. Ommatidia were gently dissociated using insect needles under a stereo microscope (600 lux). The dissection and dissociation were completed in 3 min. The preparation was immediately imaged via the Olympus AX70 microscope.

### Fluorescent Image Analyses

All image manipulations were performed under the guidelines of Rossner and Yamada (24). Fluorescent images included in the Figures are similar in appearance to the raw images. Image analyses were performed via the ImageJ program (NIH).

### Time-lapse Video Analysis

Time-lapse videos were created using ImageJ (National Institute of Health). All images acquired from fluorescence microscopy were saved in a single folder. Images were opened using ImageJ and the images were then imported as an image sequence. The image sequence was saved as an AVI file that could be viewed as a time-lapse video.

### Western Blotting

Western blotting using specific antibodies was used to confirm various mutations. Briefly, fly heads were dissected and proteins were extracted with 2X Laemmli sample buffer. Total fly proteins were size-fractionated by SDS/PAGE (10%) and transferred onto a nitrocellulose filter. Filters were incubated first with the desired primary antibodies, then with fluorophore-conjugated secondary antibodies (e.g. Alexa Fluor 680 Goat Anti-Rabbit IgG, Invitrogen). The fluorescent signal of the secondary antibodies was quantified by the Odyssey Infrared Imaging system (LI-COR, Lincoln, NE, USA). Polyclonal antibodies against Arr1, Arr2, eye-PKC, NinaC, and NorpA were used.

### Construction of rdgC-mCherry cDNA for the generation of transgenic lines

The *rdgC* gene encodes three RdgC isoforms (746, 677, and 661 aa) with varying N-terminal sequences (25). We chose the longest protein isoform as the reporter, which was incorporated with the mCherry tag at its C-terminus. Specifically, the full-length cDNA encoding the long form of RdgC (746 aa) was constructed by combining the coding sequence of a partial *rdgC* cDNA (RH46370, DGRC Stock 10792; https://dgrc.bio.indiana.edu//stock/10792 ; RRID:DGRC_10792, *Drosophila* Genetic Research Center) with a synthetic DNA sequence corresponding to the missing 5’ sequence (GenScript). The cDNA corresponding to the mCherry sequence was ligated to the 3’ end of the *rdgC* cDNA after the removal of the stop codon. The resulting *rdgC-mC* recombinant DNA was subcloned into YC4 for generating transgenic flies.

## RESULTS

### Endocytosis of Arr1-GFP and Rh1-mC upon blue light stimulation in pupal photoreceptors

It has been shown that *Drosophila* rhodopsin Rh1 displays light-dependent endocytosis in pupal photoreceptors (26). Specifically, endocytosis is regulated by the light-dependent phosphorylation of Rh1 at its C-terminus. Moreover, phosphorylation is also critical for the recruitment of the cytosolic Arr1 to the rhabdomere (26). To further explore the role of Arr1, we employed transgenic flies expressing Arr1-GFP for insights into the mechanism of endocytosis in live photoreceptors (16).

As shown previously, a continued stimulation with an intense blue light (460-500 nm, 1300 lux) resulted in the translocation of Arr1-GFP to the rhabdomere (**Fig 1** **A**), which reached a steady state in about two and half minutes (16). Subsequently, Arr1-GFP became internalized leading to its sequestration in cytoplasmic vesicles (**Fig 1** **B**); internalization reached a steady state in approximately four minutes following translocation (16). Consistently, we show Arr1-GFP was present in both rhabdomeres and cytoplasmic vesicles of dissociated photoreceptors following the light exposure (**Fig 1** **C**).

**Fig 1.**
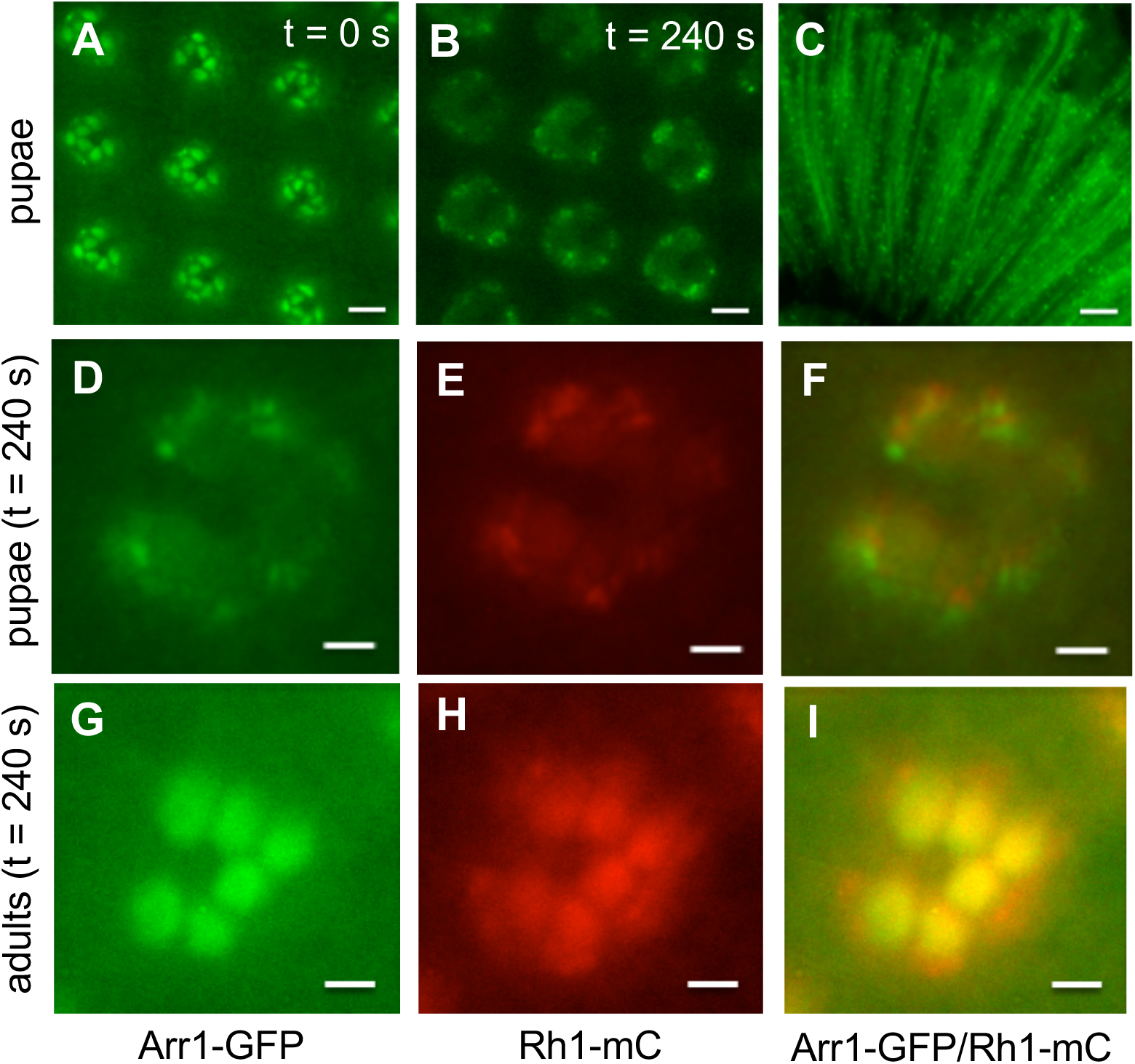
The light-dependent co-internalization of Arr1-GFP with Rh1-mC occurs in pupal but not adult photoreceptors. (A, B) Blue light triggers translocation and subsequent internalization of Arr1-GFP. Shown are the subcellular localization of Arr1-GFP at two time points following its translocation to the rhabdomere (A, t = 0 s; B, t = 240 s). (C) Arr1-GFP was observed in cytoplasmic vesicles and rhabdomeres of dissociated photoreceptors after internalization. (D, E, F) Arr1-GFP and Rh1-mC were prominently co-localized in the cytoplasmic vesicles of pupal photoreceptors following blue light stimulation. (G, H, I) Arr1-GFP and Rh1-mC were co-localized mostly in the rhabdomere of adult photoreceptors following blue light stimulation. Scale bar, 5 μm (A, B), 10 μm (C), 2 μm (D-I)

To support that Arr1 is co-internalized with Rh1, we investigated their subcellular localization. In pupal photoreceptors following the blue light stimulation, we observed that most of Arr1-GFP (**Fig 1** **D**) and Rh1-mC (**Fig 1** **E**) appeared co-localized in the vesicles within the cytoplasm as well as in the rhabdomere (**Fig 1** **F**). In contrast, in adult photoreceptors, Arr1-GFP (**Fig 1** **G**) and Rh1-mC (**Fig 1** **H**) were co-localized mostly in the rhabdomere (**Fig 1** **I**) as Arr1-GFP failed to internalize and thus not co-localized with Rh1-mC in the cytoplasm (16).

### Endocytosis of Arr1-GFP is blocked in the ninaC mutant affecting the unconventional myosin III

NinaC encodes two myosin III polypeptides, p132 and p174 (27), which are localized in the cytoplasm or rhabdomeres, respectively (28). The NinaC proteins have been shown to play a role in the transport of several visual signaling proteins (29–31). We investigated how NinaC might be involved in the light-dependent translocation and/or endocytosis of Arr1-GFP. Interestingly, we observed that the light-dependent endocytosis of Arr1-GFP was blocked in the *ninaC^3^* (*ninaC^P225^*) mutant (**Fig 2** **B-D).** However, the translocation of Arr1-GFP to the rhabdomere was not affected. The *ninaC^3^*allele lacks the p174 isoform due to a stop codon deleting the last 440 aa from the C-terminus of p174 while p132 is present but missing the last 54 aa (27). Based on the findings, it appears that NinaC plays a role in modulating the endocytosis of Arr1-GFP.

**Fig 2.**
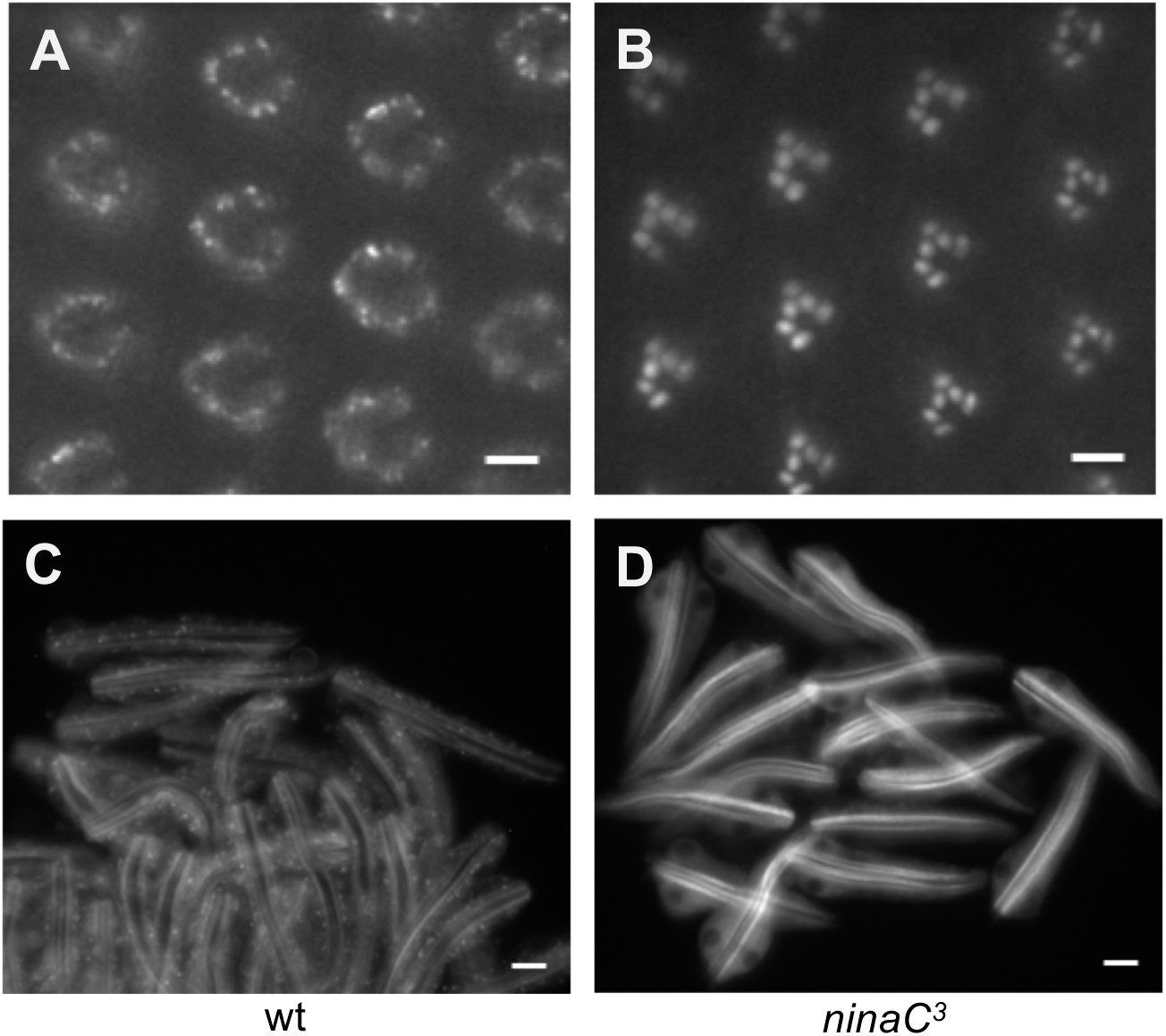
The light-dependent internalization of Arr1-GFP is eliminated in *ninaC^3^* mutants. (A) Arr1-GFP displayed endocytosis in wild-type but not in *ninaC^3^*(B) background following blue light illumination for five minutes. Shown are the subcellular localization of Arr1-GFP (A, B) in the eye and in dissociated photoreceptors (C, D) after light stimulation. Scale bar, 5 μm (A, B), 10 μm (C, D)

### Internalization of Arr1-GFP is regulated by dynamin and is eliminated in mutants affecting the biosynthesis of PIP_2_

We further explored whether CME (22) is responsible for the endocytic trafficking of Arr1-GFP by examining the contribution of dynamin, a GTPase involved in the scission of vesicles (32). Dynamin is encoded by the *shibire* (*shi*) gene (19, 33) in *Drosophila.* Therefore, we investigated whether the endocytosis could be affected in the temperature-sensitive *shi^ts1^* background (19). Importantly, we observed that internalization of Arr1-GFP was eliminated when mutant flies were analyzed at the restrictive temperature (>29° C) (**Fig 3** **B**) in which dynamin becomes inactivated. In contrast, endocytosis was not affected when flies were analyzed at the permissive temperature (20°C) (**Fig 3** **C**). Together, our findings strongly support that the blue light-initiated endocytosis of Arr1-GFP is dependent on dynamin.

**Fig 3.**
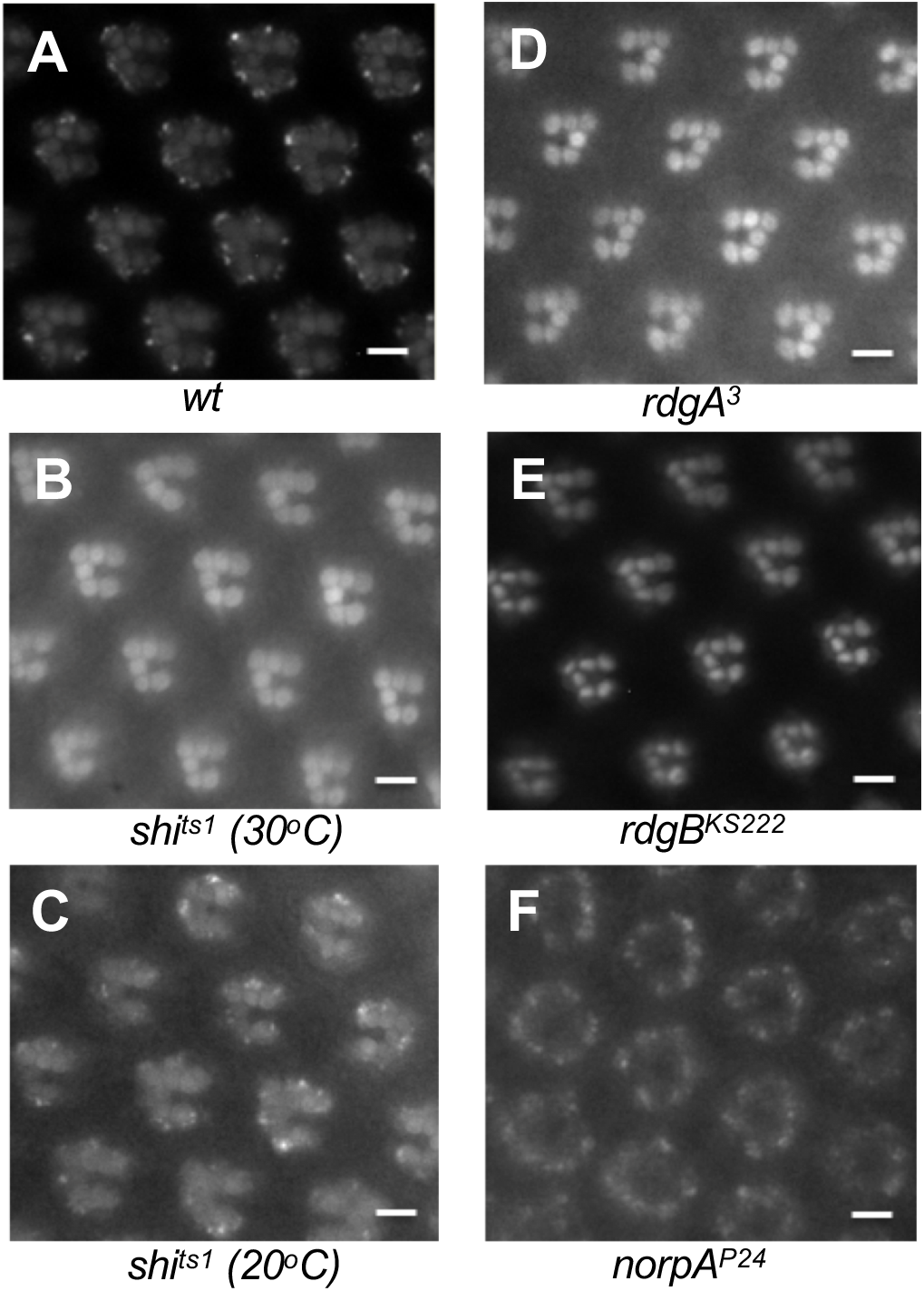
The blue-light initiated endocytosis of Arr1-GFP is blocked in *shi^ts1^*, *rdgA^3^*, or *rdgB^ks222^*, but not in *norpA^P24^* mutants. (A) In wild-type photoreceptors Arr1-GFP displayed light-dependent internalization. (B) Internalization was eliminated in *shi^ts1^* mutants at the restrictive temperature (30° C) but not at the permissive temperature (20° C) (C). (D, E) The internalization was also blocked in either *rdgA^3^* (D) or *rdgB^ks222^* mutants (E). (F) The internalization was observed in *norpA^P24^* photoreceptors. Images of the compound eye were taken after about five minutes of the blue light exposure. Scale bar, 5 μm

CME also is regulated by phosphatidylinositol 4, 5-bisphosphate (PIP_2_) (34), a minor phospholipid in the membrane. We explored whether a perturbation of the PIP_2_ biosynthesis impacted the endocytosis of Arr1-GFP. The biosynthesis and recycling of PIP_2_ require a cascade of biochemical reactions that convert diacylglycerol (DAG) to PIP_2_, which include RdgA, a DAG kinase that catalyzes the conversion of DAG to phosphatidic acid (20), and RdgB, a phosphatidylinositol transfer protein (21). Significantly, we show that endocytosis of Arr1-GFP was blocked in either *rdgA^3^*(**Fig 3** **D**) or *rdgB^KS222^* mutants (**Fig 3** **E**). In contrast, endocytosis was not affected by the *norpA* mutation (**Fig 3** **F**) that lacks phospholipase Cβ involved in the light-dependent hydrolysis of PIP_2_ (35). Taken together, we conclude that the blue light-mediated endocytosis of Rh1 with the co-internalization of Arr1-GFP mimicking the stimulation of an agonist to its cognate GPCR, requires both dynamin and PIP_2_ and thus likely belongs to CME (22).

### Green light triggers the endocytosis of Rh1-mC in both adult and pupal photoreceptors

Rh1 can be optimally activated by the blue light (460-500 nm) flanking its absorption maximum (17) to trigger CME. Rh1 also can be activated by the green light (530-560 nm), which may behave like a partial agonist. Green light may evoke a distinct conformation of Rh1 different from that of the blue light. We explored the endocytic mechanism of Rh1 by green light in transgenic flies expressing Rh1-mC (36).

As shown in Figure 4, Rh1-mC was initially present in the rhabdomere of R1-6 photoreceptors (**Fig 4** **A, C**). A continuous green light exposure triggered its internalization leading to the accumulation in the cytoplasmic vesicles of both adult (**Fig 4** **B)** and pupal photoreceptors (**Fig 4, D-E, S1 Fig**). These cytoplasmic vesicles appeared to form larger vesicles similar to RLVs or Rh1-immunopositive large vesicles (26), which were present surrounding the base of rhabdomeres (**Fig 4** **F**). In pupal photoreceptors we show that Rh1-mC displayed a trafficking trajectory similar to that of Arr1-GFP but with faster kinetics; endocytosis of Rh1-mC reached a steady state at approximately three minutes after light stimulation (**Fig 4** **D**). In contrast, blue light triggers the translocation and the internalization of Arr1-GFP; both of which reach a steady state at approximately six and a half minutes (16). Together, green light stimulation appears to render Rh1 in a partially active conformation, which is also subjected to regulation via internalization.

**Fig 4.**
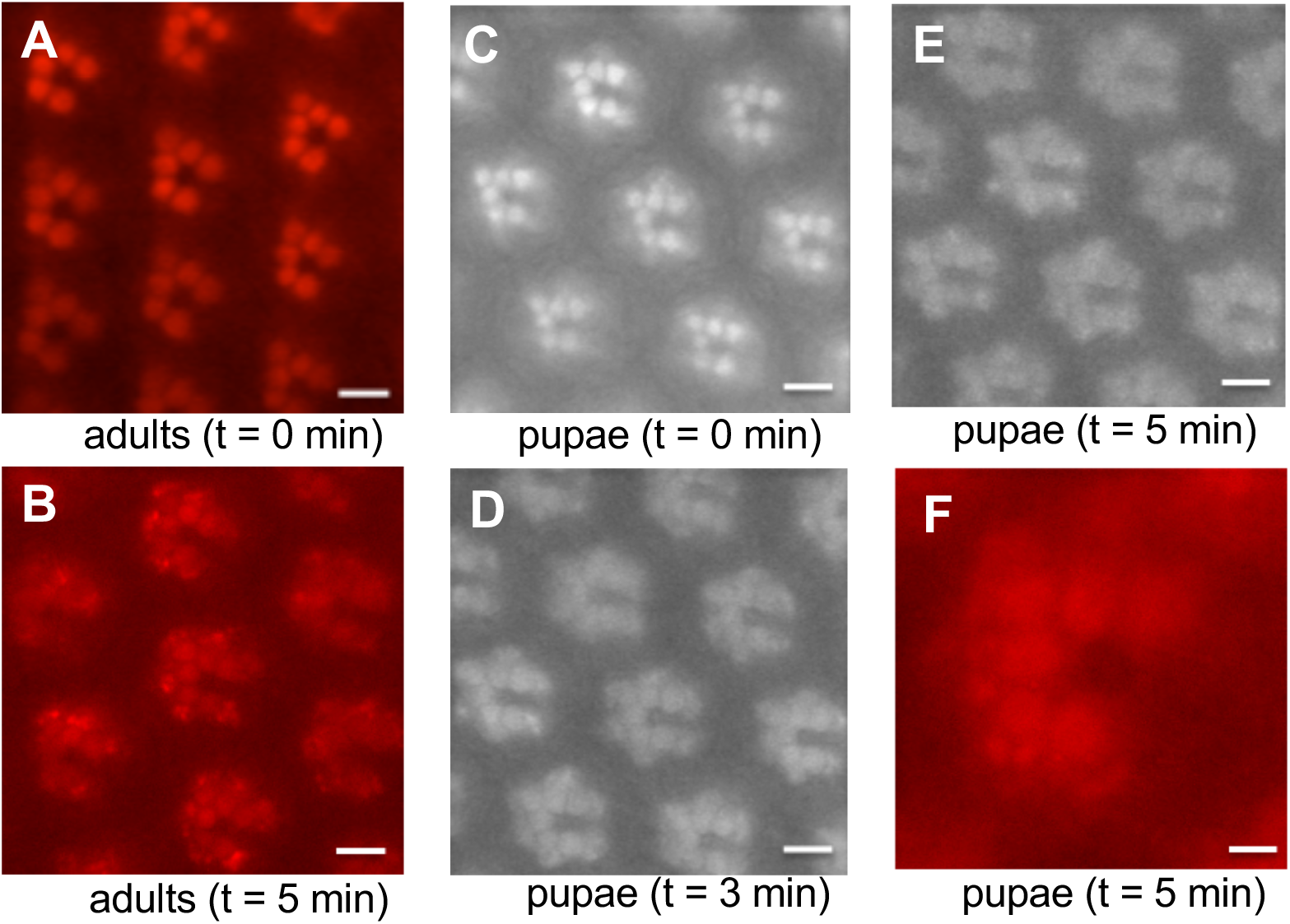
The light-dependent internalization of Rh1-mC upon green light stimulation. (A) In adult flies Rh1-mC was highly enriched in the rhabdomere of R1-6 photoreceptors of the compound eye. (B) Rh1-mC underwent endocytosis upon continued green light (530-560 nm) exposure for five minutes leading to its accumulation in the cytoplasmic vesicles. (C-E) In pupal photoreceptors green light also triggered Rh1 endocytosis similar to that observed in adult photoreceptors. (F) The internalized Rh1-mC was localized in large vesicles surrounding the rhabdomere. t, time following the light stimulation. See Supplemental Movie 1 for the green light-mediated endocytosis of Rh1-mC in pupal photoreceptors. Scale bar, 5 μm (A-E), 2 μm (F)

### Endocytosis of Rh1-mC is independent of Arr1 and Arr2

We next investigated whether Arr1 or Arr2 plays a role in orchestrating the light-dependent endocytosis of Rh1-mC in pupal photoreceptors. However, we show the endocytosis was not eliminated in either *arr1^1^* (**Fig 5, C-D**) or *arr2^3^* mutants (data not shown), suggesting that neither Arr1 nor Arr2 serves as the adaptor protein in promoting the internalization.

**Fig 5.**
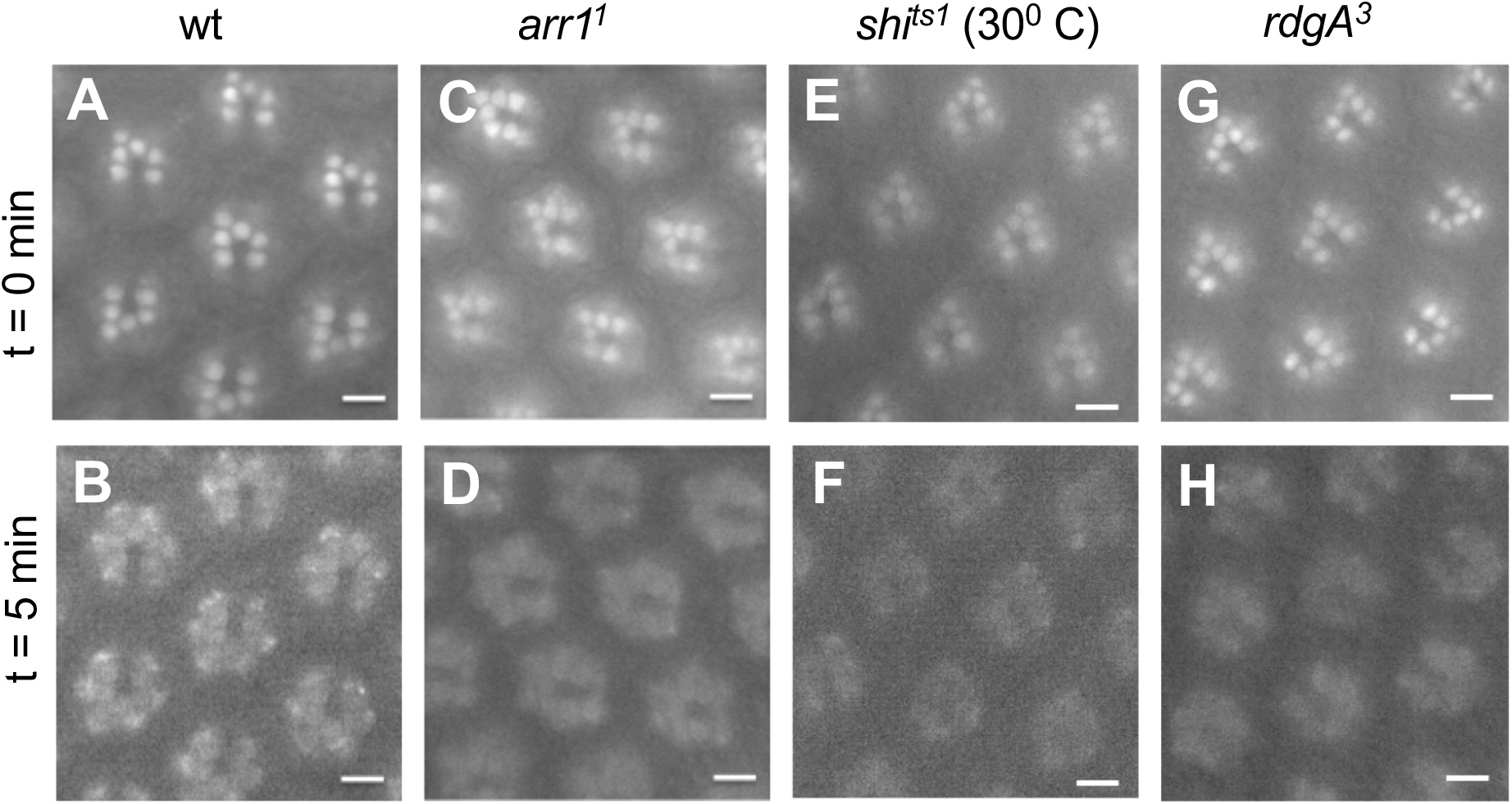
Endocytosis of Rh1-mC is not affected in *arr1^1^, shi^ts1^*, or *rdgA^3^* mutants. The green light-mediated internalization of Rh1-mC was not affected in *arr1^1^* (D), *shi^ts1^* (at 30°C) (F), or *rdgA^3^* (H) mutants. Fluorescent images were taken initially (t = 0 min, top panel) and after five minutes of light exposure (t = 5 min, bottom panel). Scale bar, 5 μm

The lack of involvement by Arr1 might be attributed to the lack of phosphorylation in Rh1. As phosphorylation of Rh1 is required for the Arr1 association, we explored whether Arr1 could be recruited by Rh1 following the green light stimulation. Indeed, we show green light exposure led to the translocation of over 50% of Arr1-GFP to the rhabdomere (**Fig 6** **B**), indicating that Rh1 was phosphorylated. Furthermore, translocation of Arr1 appeared accompanied by the endocytosis of Rh1-mC (**Fig 6** **A**). In contrast, blue light stimulation triggered the internalization of both Rh1-mC (**Fig 6** **D**) and Arr1-GFP (**Fig 6** **E**); both of which were prominently co-localized in the cytoplasmic vesicles of pupal photoreceptors (**Fig 6** **F**).

**Fig 6.**
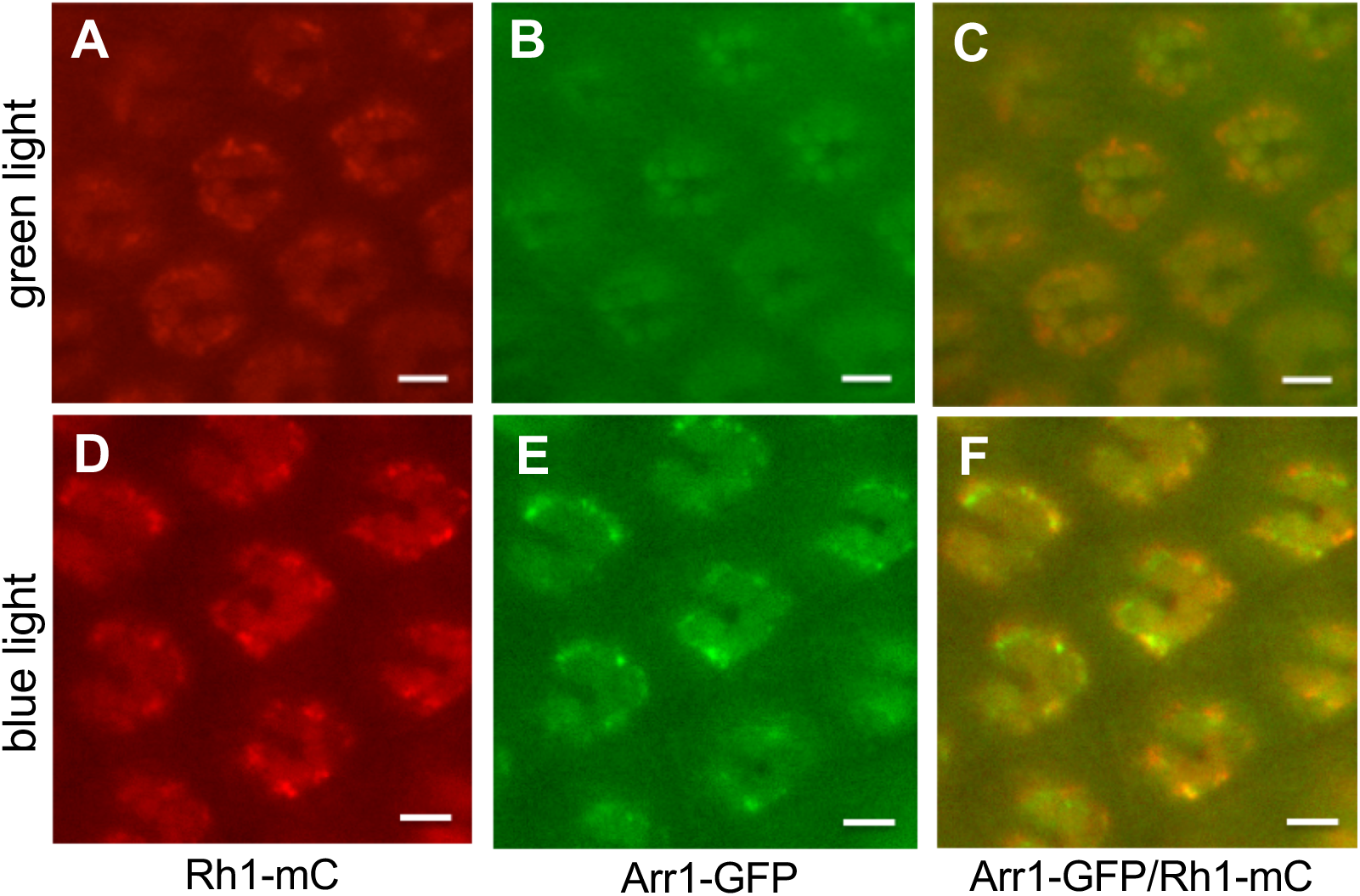
The subcellular distribution of Rh1-mC and Arr1-GFP following either green or blue light stimulation of pupal photoreceptors. (Top panel) Green light triggered endocytosis of Rh1-mC (A), which is accompanied by recruitment of over 50% of Arr1-GFP to the rhabdomere (B). Rh1-mC and Arr1-GFP appeared mostly co-localized in the rhabdomere (C). (Bottom panel) Blue light triggered endocytosis of both Rh1-mC (D) and Arr1-GFP (E) and both were co-localized in cytoplasmic vesicles and the rhabdomere (F). Scale bar, 5 μm

Taken together, the green light exposure also leads to phosphorylation of Rh1 that recruits Arr1 to the rhabdomere. However, this interaction may be transient as Arr1 appears not directly involved in the subsequent endocytic event.

### Endocytosis of Rh1-mC in pupal photoreceptors is not blocked in shi nor rdgA mutants

We explored whether the light-dependent endocytosis of Rh1-mC also involves dynamin. Therefore, we tested in the *shi^ts1^* mutant background at the restrictive temperature (30^0^ C) (19, 33) and observed the internalization was not affected (**Fig 5, E-F**). We also investigated whether a reduction of the PIP_2_ level due to a lack of the RdgA protein impacts endocytosis (20, 37). However, it was not perturbed in the *rdgA^3^* mutant background (**Fig 5, G-H**). Based on the findings, we conclude that the green light-mediated internalization of Rh1-mC does not belong to CME as it is independent of dynamin and PIP_2_.

Taken together, activated rhodopsin can be subjected to two distinct endocytic trafficking mechanisms, a dynamin-dependent, and a dynamin-independent mechanism, depending on the nature of light stimulation. The blue light stimulation may mimic the effect of an agonist that promotes a stable association between Rh1 and Arr1 leading to CME while the green light might mimic a partial agonist resulting in a transient interaction that initiates the dynamin-independent endocytic mechanism.

### The fate of the internalized Rh1-mC following the light-dependent endocytosis

We explored the post-endocytic trafficking of Rh1 following these two distinct endocytic events. We first investigated if a transient interaction between Rh1 and Arr1 would readily promote the recycling of the internalized Rh1. Experimentally, the green light was used to initiate the endocytosis of Rh1-mC (**Fig 7** **A, D**); following different periods of dark adaptation photoreceptors were re-examined for subcellular distribution. Interestingly, we show that Rh1-mC could be detected in the rhabdomere after dark adaptation for two min (**Fig 7** **B**), and after five min up to 95% of the endocytosed Rh1-mC appeared recycled back to the rhabdomere (**Fig 7** **E**), when compared to prior to the light exposure (**Fig 7** **C**). Taken together, the internalized Rh1 from the non-CME appears readily recycled and re-inserted into the rhabdomere membrane.

**Fig 7.**
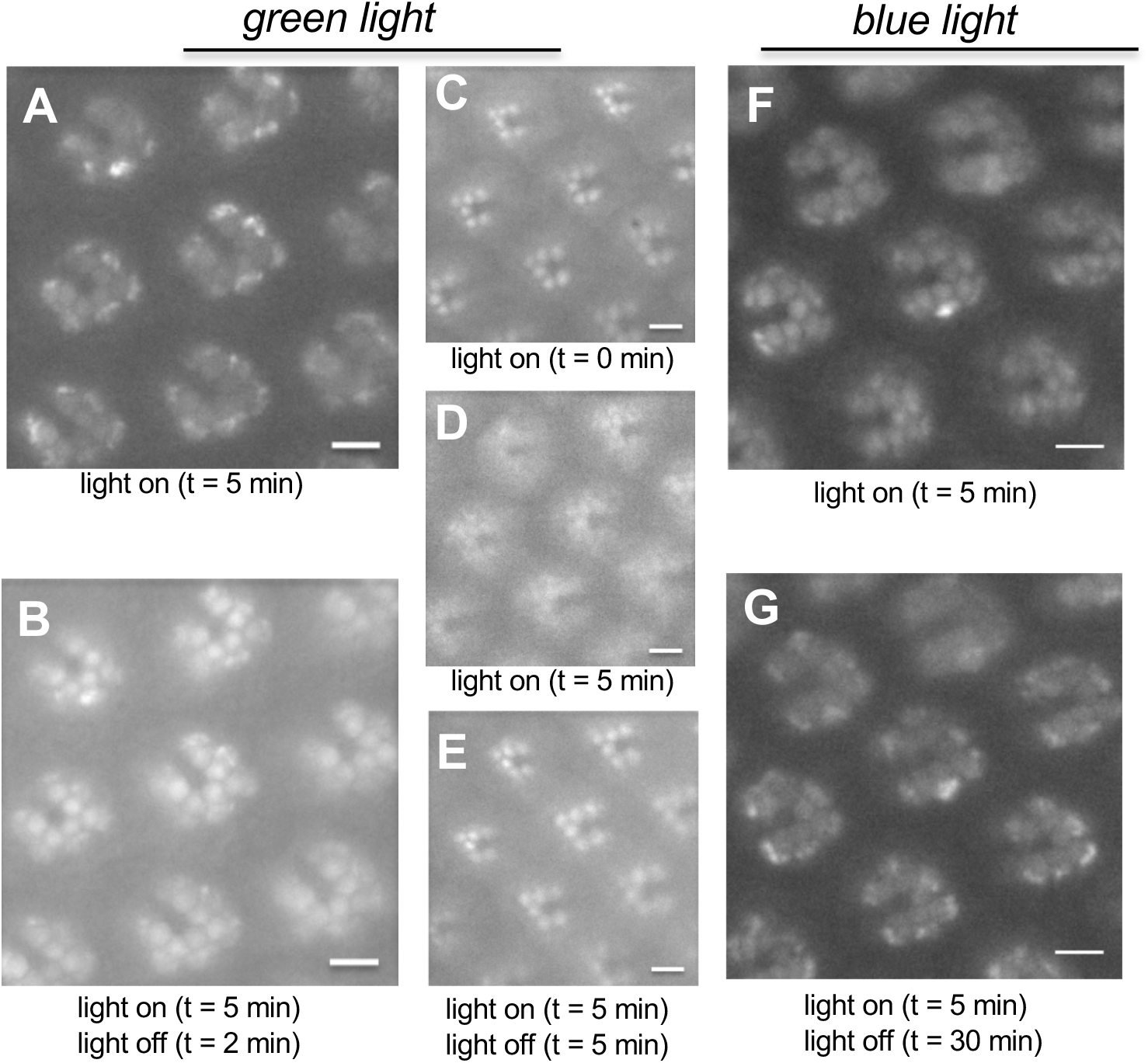
Recycling of the internalized Rh1-mC in pupal photoreceptors following dark adaptation. (A-E) Recycling of the internalized Rh1-mC from non-CME via green light stimulation. Shown are the subcellular distribution of the internalized Rh1-mC after light exposure for 5 min (A, D) and following dark adaptation for two min (B) or five min (E). (F, G) Internalized Rh1-mC from blue light-initiated CME appears not recycled following dark adaptation. Shown are the subcellular localization of Rh1-mC after light exposure for 5 min (F) and following dark treatment for 30 min (G). Scale bar, 5 μm

We also examined the fate of the internalized Rh1-mC after the blue light-mediated CME. However, we observed that the internalized Rh1mC remained in the cytoplasmic vesicles even after dark treatment for 30 min (**Fig 7, F-G**). Thus, it appears that Rh1 is not promptly recycled for re-insertion into the rhabdomere following CME. Together, the endocytic trafficking and recycling of activated rhodopsin may be somehow coordinated as rhodopsin might be sorted early to organize both the internalization and post-endocytic mechanisms.

### Differential dephosphorylation of activated rhodopsin leads to distinct post-endocytic trafficking

Recycling of the internalized Rh1 may require dephosphorylation before its trafficking back to the rhabdomere membrane. Dephosphorylation of Rh1 involves rhodopsin phosphatase encoded by the *rdgC* gene (38, 39). We investigated where dephosphorylation took place following internalization using a modified RdgC with a fluorescent mCherry tag (RdgC-mC) as the reporter.

We generated and characterized the transgenic flies expressing RdgC-mC by examining their distribution in photoreceptors. We show RdgC-mC was localized surrounding the rhabdomere distinct from Arr2-GFP present in the rhabdomere. (**Fig 8, A-C**), when visualized via deep pseudopupil (dpp). Dpp is the superimposition of virtual images from several adjacent ommatidia and it reveals the trapezoid arrangement of rhabdomeres in ommatidia. Significantly, RdgC-mC translocated to the peri-rhabdomeric area following continuous light stimulation (**Fig 8** **E**) as it appeared co-localized with internalized rhodopsin in adult photoreceptors. Indeed, the translocation of RdgC-mC reached a steady state in about three minutes, similar to the kinetics of the light-induced internalization of Rh1-mC (**Fig 4**). Importantly, translocation was eliminated in mutants with a low Rh1 content (e.g. *ninaE*) (data not shown), supporting that the interaction with phosphorylated Rh1 is responsible for the light-dependent translocation.

**Fig 8.**
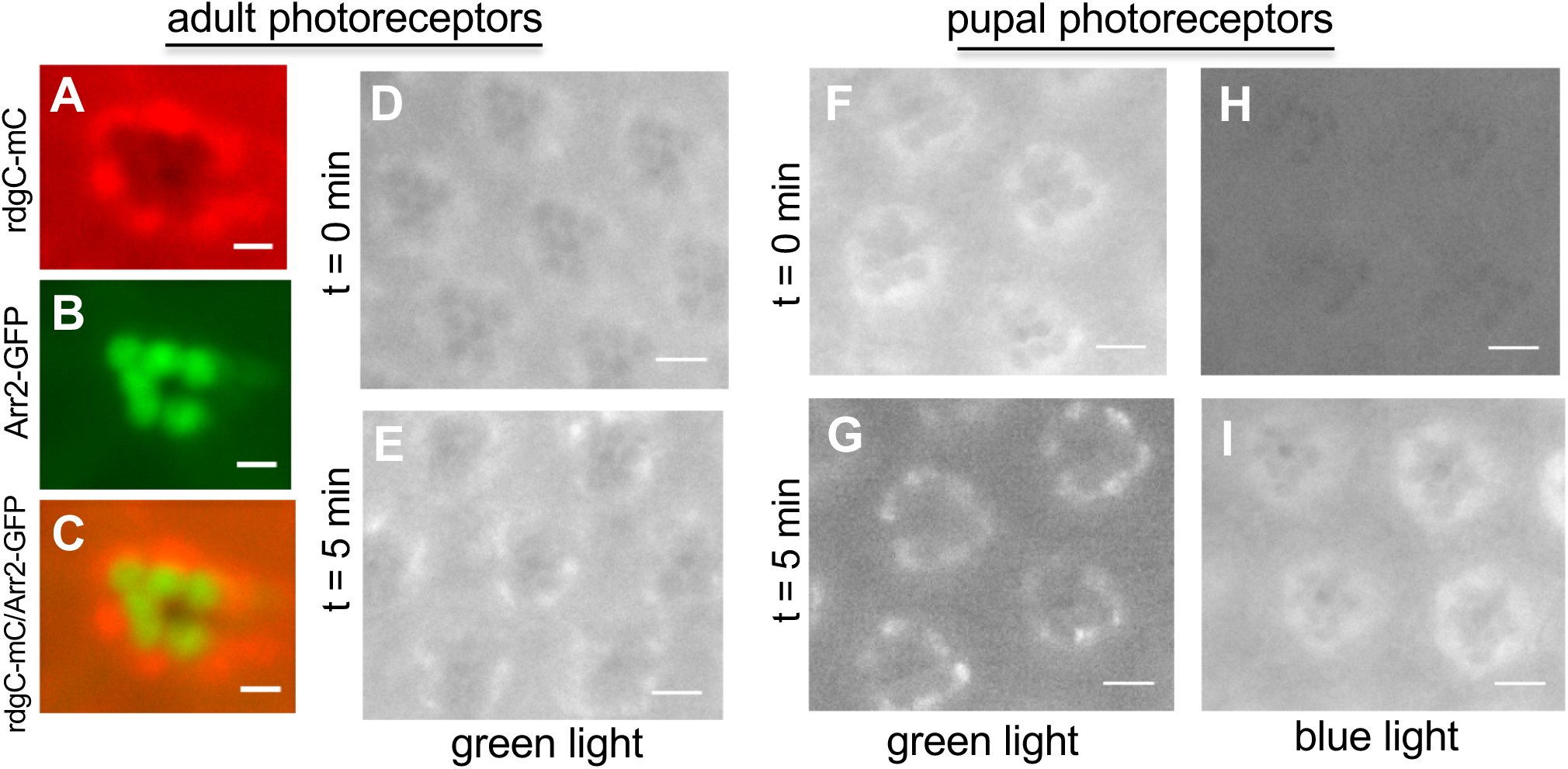
Exploring the subcellular distribution of phosphorylated Rh1 in post-endocytic trafficking via RdgC-mC. (A) RdgC-mC was localized in the cytoplasm of adult photoreceptors. Shown is the distribution of RdgC-mC as visualized in dpp (deep pseudopupil) of the compound eye. (B) Arr2-GFP was localized in the rhabdomere due to its association with activated Rh1. (C) RdgC-mC and Arr2-GFP displayed distinct subcellular localization in the eye as shown in the merged images. (D) RdgC-mC appeared uniformly distributed in the cytoplasm surrounding the rhabdomere of photoreceptors at the beginning of light stimulation. (E) RdgC-mC translocated towards the base of rhabdomeres following green light stimulation for five min. (F-G) Green light stimulation of pupal photoreceptors led to the enrichment of RdgC-mC at the base of the rhabdomere. (H-I) Blue light stimulation for five min exerted no effect on the enrichment as RdgC-mC remained in the cytoplasm. (H) Shown are the rhabdomeres of ommatidial clusters when stimulated with blue light in RdgC-mC expressing flies. Images were taken at t = 0 min or t = 5 min after the initiation of the light. Scale bar, 10 μm (A-C), 5 μm (D-I)

We compared the subcellular distribution of RdgC-mC following the endocytosis of Rh1 in pupal photoreceptors. In non-CME, we show that RdgC-mC became enriched at the base of rhabdomeres (**Fig 8** **G**), suggesting that it was recruited to the internalized Rh1 for dephosphorylation. Timely dephosphorylation of phosphorylated Rh1 by RdgC may be critical for the recycling of Rh1 following the non-CME (**Fig 7C, E**). In contrast, following the blue light stimulation RdgC-mC failed to translocate towards the base of rhabdomeres (**Fig 8, I**). Instead, it appeared uniformly distributed in the cytoplasm of photoreceptors, suggesting that RdgC-mC was incapable of recognizing phosphorylated Rh1 following CME. The lack of interaction might be due to the fact that phosphorylated Rh1 formed a stable complex with Arr1 that prevents the interaction.

Based on the findings, we conclude that differential dephosphorylation of internalized Rh1 plays a critical role in the resensitization and post-endocytic recycling processes. Moreover, Arr1 negatively regulates dephosphorylation of Rh1 by preventing the recruitment of RdgC leading to a delay of the recycling of phosphorylated Rh1 in CME.

## DISCUSSION

In this report we employed the use of *Drosophila* Rh1, taking advantage of the fact that Rh1 can be activated by different wavelengths of light to generate distinct conformations, for insights into mechanisms of the endocytic and post-endocytic trafficking in vivo. Specifically, Rh1 can be optimally activated by blue light (460-500 nm) similar to the use of a full agonist. Indeed, blue light triggers endocytosis of Rh1-mC and Arr1-GFP, which appears to belong to CME as it is dependent on dynamin and is regulated by the PIP_2_ content of the membrane. In contrast, green light (530-560 nm) stimulation which probably functions like a partial agonist, triggers the endocytosis of Rh1-mC but not Arr1-GFP. This endocytosis is also distinct from CME as it does not require dynamin and is independent of PIP_2_. Moreover, it is not regulated by Arr1, although Arr1 is recruited to the rhabdomere. Our study demonstrates differential activation of Rh1 leads to two distinct endocytic mechanisms. We also show the endocytosis and post-endocytic mechanisms appear coordinated. Furthermore, Arr1 negatively affects the recycling of the internalized Rh1 by preventing dephosphorylation. We propose that differential activation of GPCR determines the phosphorylation patterns that regulate the interaction with arrestin to impact both the endocytic trafficking and recycling of the receptor *in vivo*.

Our genetic analyses support that the blue-light mediated endocytosis of Rh1 and Arr1 belongs to CME. CME is involved in most of the agonist-induced internalization of GPCRs, in which arrestin serves as the adaptor protein by tethering activated GPCRs to the clathrin-coated pits. CME is regulated by PIP_2_, which is required by several endocytic proteins including dynamin (34). We show that the blue light-initiated co-endocytosis of Arr1-GFP is blocked in the *rdgA^3^* or *rdgB^KS222^* mutants. Both RdgA (20, 37) and RdgB (21, 40) participate in the biosynthesis and regeneration of PIP_2_ in the membrane. CME also requires dynamin, a GTPase involved in membrane scission of the vesicle (32). Consistently, internalization of Arr1-GFP is abolished in the *shi*^ts1^ mutant at the restrictive temperature, which expresses a temperature-sensitive dynamin with a point mutation at 268 aa (19). In contrast, the green light-mediated endocytosis of Rh1-mC is independent of dynamin and insensitive to a reduction of PIP_2_. Based on the findings, it appears that the light-dependent endocytosis of Rh1 is regulated by its differential phosphorylation that modulates its interaction with Arr1 to orchestrate distinct internalization events. The nature of the dynamin-independent endocytosis of Rh1 remains to be explored.

### Regulation of CME by NinaC

Our study uncovered a novel function of NinaC for promoting the endocytosis of Rh1 by the dynamin-dependent CME. The NinaC gene encodes two isoforms of myosin III (27), designated as p132 and p174 with distinct subcellular localizations. The p132 polypeptide is present in the cytoplasm while p174 is associated with the rhabdomere of photoreceptors. NinaC also contains an EF-hand domain (27) and is the major calmodulin-binding protein in photoreceptors (41). Loss of p174 leads to slower termination of the visual signaling and retinal degeneration (42). Interestingly, NinaC has been implicated in the transports of several visual signaling proteins including Arr2, Gqα, and Trpl. Specifically, Trpl, the cation channel involved in the light-dependent depolarization (43), is trafficking out of rhabdomeres and sequestrated in the cytoplasmic compartment following light stimulation, similar to endocytosis. However, the trafficking of Trpl is independent of dynamin (30). In contrast, the α-subunit of Gq undergoes light-dependent translocation by trafficking to the rhabdomere (29) from the cytoplasm. Similarly, Arr2 also translocates to the rhabdomere following the activation of the visual signaling (31). Translocation of both Arr2 and Gq is reduced in the *ninaC* mutants (29, 31). How does NinaC regulate the endocytosis of Rh1 in CME remains to be explored.

### Phosphorylation of GPCRs and regulation of internalization

It is well established that agonist-activated GPCRs are phosphorylated at multiple residues at the C-terminal sequence and/or the third cytoplasmic loop of the receptor in a ligand- and GRK-dependent manner (44). In general, phosphorylation occurs sequentially or in a hierarchical manner (45). Moreover, the order of phosphorylation appears dependent on accessibility as residues close to the C-terminus are more readily phosphorylated (46).

The extent of GPCR phosphorylation may correlate with the propensity of the phosphorylated receptor to interact with arrestin which initiates the internalization. It appears that the more phosphorylated residues in the receptor the stronger the affinity for arrestin, as demonstrated in the mu-opiate receptor (MOR) (47, 48). For example, a potent agonist D-Ala2-MePhe4-Glyol5 Enkephalin (DAMGO) triggers phosphorylation at Ser^375^, Thr^370^, Thr^376^, and Thr^379^ leading to internalization of the receptor. In contrast, morphine, a weak agonist, results in limited phosphorylation at Ser^375^, which leads to a transient arrestin interaction that is insufficient to drive the internalization. A transient arrestin interaction was observed when isoproterenol was used to activate the β-adrenergic receptor leading to endocytosis. However, the internalized β-receptor is readily recycled, suggesting that a weak arrestin interaction also promotes recycling of the receptor (49). Recycling of GPCRs may also involve a diverse set of C-terminal sequences in the receptor including the PDZ ligand (50, 51).

In *Drosophila*, Rh1 is known to undergo the light-dependent phosphorylation that is required for the Arr1 interaction (26). Phosphorylation of Rh1 takes place at the C-terminus as a truncation mutant (Rh^Δ356^) that lacks the last 17 aa eliminates the phosphorylation *in vivo* (39). Within the last 17 aa, ^357^SSDAQSQATASEAESKA^373^, there are six putative phosphorylation sites. Substitutions of Ser/Thr with Ala/Val in all six residues in flies expressing Rh1CT S>A prevented the Rh1 internalization and Arr1 translocation *in vivo* (26), supporting the critical role of the C-terminal phosphorylation. It remains to be investigated which residues of Rh1 are phosphorylated *in vivo* and what is the order of phosphorylation under different light conditions.

### The phosphorylation barcode hypothesis

It has been proposed that agonist-induced phosphorylation of GPCRs generates diverse phosphorylation patterns that are critical for the downstream signaling events. The different phosphorylation signatures are mostly contributed by GRKs as each kinase displays distinct tissue-specific expression and substrate specificity (“barcoding”) (52, 53). Subsequently, phosphorylated receptors may interact with specific conformations of arrestin to initiate selective biological effects including desensitization, endocytosis, and alternate signaling. Here we present our *in vivo* evidence related to the barcoding of GPCRs in which differential phosphorylation states of rhodopsin induced by different wavelengths of light lead to two distinct endocytic and post-endocytic trafficking, which is contributed in part by differential interactions with Arr1.

### Differential regulation of endocytosis and recycling of rhodopsin by Arr1 in *Drosophila*

We propose that green light elicits a conformation of Rh1 leading to perhaps a more ‘limited’ phosphorylation at its C-terminus. Consequently, phosphorylated Rh1 displays a transient interaction with Arr1 promoting its translocation to the rhabdomere but not co-internalization. Therefore, without Arr1 the internalized Rh1 readily associates with RdgC for dephosphorylation in the endocytic compartment leading to fast recycling (in the non-CME, **Fig 9**). In contrast, the blue light stimulation may result in more extensive phosphorylation of Rh1 leading to a stable Arr1 interaction that promotes the internalization of the Rh1-Arr1 complex. Importantly, the internalized Arr1 appears to impede the recruitment of RdgC delaying dephosphorylation of internalized Rh1 for recycling (in CME, **Fig 9**).

**Fig 9.**
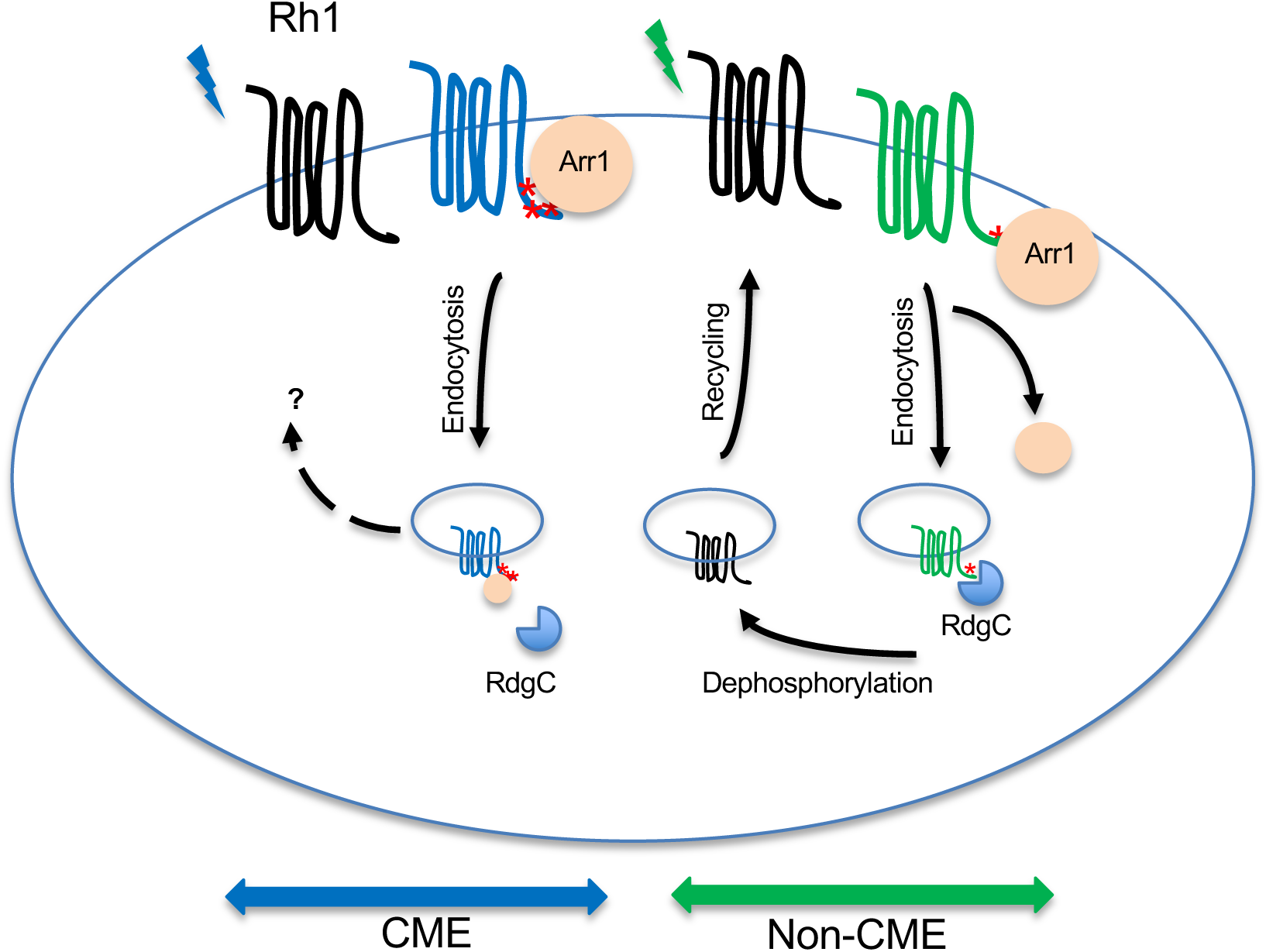
The two distinct endocytic and post-endocytic trafficking mechanisms of activated Rh1 in pupal photoreceptors. (Left), In the blue light-initiated CME, multi-phosphorylated Rh1 forms a stable complex with Arr1 leading to its co-internalization. Subsequently, Arr1 may interfere with the recruitment of RdgC that dephosphorylates Rh1 delaying the recycling of Rh1. Right, In the dynamin-independent non-CME, phosphorylated Rh1 is internalized without Arr1 enabling timely dephosphorylation by RdgC to promote recycling. Phosphorylated residues at the C-terminus of Rh1 are indicated as red *.

To sum up, our findings support the notion that differentially activated Rh1 influences the extent of its phosphorylation that controls both trafficking and recycling mechanisms via differential interactions with Arr1. Arr1 plays a major role in coordinating the desensitization and resensitization of Rh1 by controlling the interaction with clathrin-coated pits and rhodopsin phosphatase, respectively.

## Supporting information

The light-dependent endocytosis of Rh1-mC

## ACKNOWLEDGEMENTS

We thank the Drosophila Genomics Resource Center for the *rdgC* cDNA clone (NIH Grant 2P40OD010949). We thank technical help from Jorge Alejandro Antunez, Joshua Lee, Bryan Mainhardt, and Carolyn Yee. This work was supported by NIH grants [R01, to B-H S].

## Abbreviations

Arr1: arrestin 1
Arr2: arrestin 2
CME: clathrin-mediated endocytosis
DAG: diacylglycerol
dpp: deep pseudopupil
GFP: green fluorescent protein
GPCRs: G-protein coupled receptors
GRK: G-protein-coupled receptor kinase
ninaC: neither-inactivation-no-afterpotential C
PIP_2_: phosphatidylinositol 4, 5-bisphosphate
rdgA: retinal degeneration A
rdgB: retinal degeneration B
rdgC: retinal degeneration C
Rh1: rhodopsin 1
shi: shiberi

## SUPPORTING INFORMATION

**Supplemental Movie 1. Light-dependent endocytosis of Rh1-mCherry in pupal photoreceptors.** Time-lapse images were obtained by fluorescence microscopy and processed to generate a movie using ImageJ. Images were taken every 20 seconds for 4 minutes

## Notes

### Competing Interest Statement

The authors have declared no competing interest.

